# Interventional breath analysis reveals oral microbial conversion of 1,2-Propanediol into Propanal and 1-Propanol

**DOI:** 10.64898/2026.11.05.764750

**Authors:** Anesu Chawaguta, Daniel Sanders, Lorenzo S Petralia

## Abstract

**Background:** Orally exhaled breath contains volatile organic compounds (VOCs) that may reflect local microbial metabolism as well as systemic processes. Consequently, the breath sampling route can influence VOC profiles and complicate biomarker interpretation. Previous studies have shown that 1-propanol is substantially enriched in orally compared with nasally exhaled breath.

**Objective:** To determine whether a controlled oral challenge with 1,2-propanediol (1,2-PD) induces sampling route-dependent changes in breath 1-propanol concentrations.

**Methods:** Ten healthy volunteers received 0.1 mL 1,2-PD diluted in approximately 50 mL water. The solution was used as an oral rinse for approximately 30 s and then expectorated. Matched oral and nasal end-tidal breath samples were collected before administration, immediately afterwards, and at 15-min intervals. Propanal, 1-propanol, acetol, acetone, and isoprene were analysed using gas chromatograph-ion mobility spectrometer (GC-IMS).

**Results:** Oral 1,2-PD exposure produced a rapid and transient increase in exhaled propanal and 1-propanol signals, with the largest responses immediately after administration and at the second post-intervention measurement. In contrast, nasal signals remained comparatively stable. Acetone and isoprene showed no consistent intervention-related changes, regardless of the sampling route. No reproducible increase in acetol was detected, although interpretation was limited by analytical interference.

**Conclusion:** A controlled oral 1,2-PD challenge elicited a route-specific propanal and 1-propanol response, consistent with local microbial metabolism of 1,2-PD within the oral cavity. Both fucose and 1,2-PD have been previously reported in human saliva, including elevated concentrations in selected disease-associated contexts. Bacteria can degrade salivary mucin glycans, releasing sugars including fucose. Together, these observations motivate investigation of a proposed pathway linking mucin-derived fucose to 1,2-PD and subsequent propanal and 1-propanol formation in the human oral cavity.

Fig. 0.
ToC abstract figure depicting the overview of the study protocol. Image created using Biorender software.

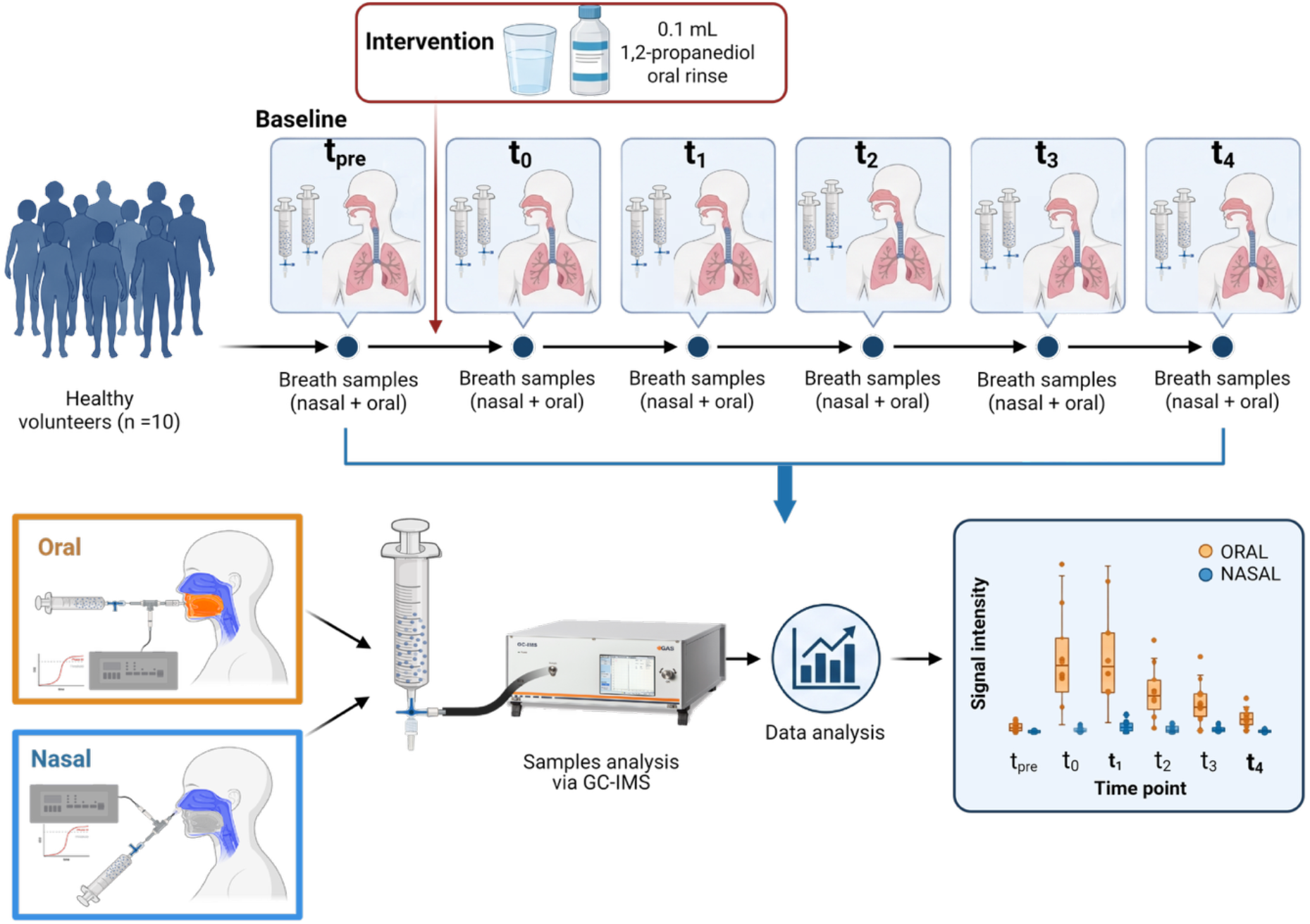

## 1 Introduction

Exhaled volatile organic compounds (VOCs) can provide a non-invasive molecular readout of metabolic, physiological, inflammatory, and pathophysiological processes occurring throughout the body. Their analysis has therefore attracted substantial interest for disease detection, monitoring, phenotyping, and precision-medicine applications. However, the VOC profile measured in exhaled breath reflects multiple sources: endogenous metabolism, respiratory and airway processes, environmental exposure, and biochemical activity within the oral cavity [1,2]. Consequently, VOCs measured in orally exhaled breath do not necessarily originate exclusively from the systemic circulation or lower respiratory tract [3]. Oral microorganisms can transform locally available substrates into volatile products, that are subsequently incorporated into mouth-exhaled breath [4,5].

This distinction has direct consequences for breath-biomarker research. A compound enriched in oral breath may reflect local microbial metabolism rather than systemic disease biology. Conversely, oral VOCs may provide functional information about the microbial ecosystem when their sources are appropriately characterised. Sampling route is therefore a critical pre-analytical variable. Controlled comparisons of oral and nasal breathing have shown that route switching can alter the measured concentrations of several VOCs, although the magnitude and direction of the effect vary between compounds [6–9]. Methodological reviews have similarly emphasised the need to control sampling conditions when breath VOCs are interpreted as biomarkers [3,10,11].

Previous route-resolved measurements identified propanol, acetoin, ethanol as strongly associated with oral sampling [6]. This association persisted under fasting conditions, reducing the likelihood that recent dietary exposure alone explained the oral enrichment. A subsequent sub-gram dose glucose oral intervention demonstrated that substrate exposure can rapidly alter fermentation-associated VOCs, including acetoin and ethanol [4]. These observations support a contribution from oral microbial metabolism but do not identify the substrate responsible for the observed route-dependent 1-propanol signal.

One candidate substrate axis involves 1,2-propanediol (1,2-PD, also known as propylene glycol), which can be converted through bacterial propanediol-utilisation reactions to propanal and subsequently to to 1-propanol or propionate. This reaction sequence is represented in the KEGG propanoate metabolism map00640, including the coenzyme B12-dependent conversion of 1,2-PD to propanal and the propanal-to-1-propanol branch [12]. The oral cavity contains host-derived substrates that could potentially support microbial metabolism between dietary exposures. Fucose and 1,2-propanediol have both been quantified in human saliva, while early experimental work showed that oral bacterial communities can degrade glycoprotein-associated sugars (including fucose) in saliva, leading to 1,2 PD production [13,14]. These observations identify a plausible salivary substrate context for 1,2-PD-associated microbial metabolism, although the endogenous source of 1,2-PD and its conversion to breath VOCs remain unresolved.

The present study was therefore designed to investigate the downstream volatile response to exogenous oral 1,2-PD. It was hypothesised that controlled oral exposure to 1,2-PD would induce an increase in propanal and 1-propanol in oral relative to nasal end-tidal exhaled breath. An exhalation route-dependent response would be consistent with local processing of 1,2-PD within the oral compartment and would provide an experimental basis for future isotope-resolved studies of endogenous oral substrate utilisation.

## 2. Methods

Ten healthy volunteers participated in a controlled oral 1,2-PD intervention study. Participants fasted for at least two hours before sampling and intervention. Participants were excluded if they reported use of electronic cigarettes or tobacco products within 2h before the study visit. Prior to baseline sampling were instructed to clear their nasal passages and to rest for a few minutes on a chair to reach a calm state [3]. For each participant, the intervention solution was prepared by diluting 0.1 mL of 1,2-propanediol (propylene glycol; CAS 57-55-6; 99.7%, supplied as food- and pharmaceutical-grade E 1520; Nanoprotect GmbH, Düsseldorf, Germany; 103.6 mg, 1.36 mmol) in 50 mL of water, giving a concentration of 27.2 mmol/L (2.07 g/L; 0.2% v/v). Participants rinsed the oral cavity with this solution for approximately 30 seconds and then expectorated it.

Matched oral and nasal end-tidal breath samples were collected before intervention (*t*_pre_), immediately after exposure (*t*_0_), and at four further time points at approximately 15-minute intervals (*t*_1_-*t*_4_, corresponding effectively to 15, 30, 45 and 60 minutes after exposure). Endtidal sampling was guided by continuous CO_2_ monitoring as described more in detail here [6]. Room air samples were measured before each participant sequence to characterise background signals. Moreover, headspace GC-IMS analysis of the 1,2-PD solution before administration detected no signals in the propanal or 1-propanol regions, excluding substrate impurity or direct instrumental response as explanations for the observed increases.

### 2.1 Breath Analysis

Samples were analysed using a BreathSpec gas chromatography-ion mobility spectrometry (GC-IMS) system (G.A.S., Dortmund, Germany). The instrument combines gas chromatographic pre-separation of volatile organic compounds (VOCs) with subsequent separation and detection by ion mobility spectrometry, providing two-dimensional separation according to GC retention time and IMS drift time.

A rapid 5-minute acquisition method was used to permit repeated measurements over the longitudinal sampling period. Chromatographic separation was performed using a 30 m polar MXT-WAX capillary column (30 m × 0.53 mm internal diameter, 0.50 µm film thickness), maintained isothermally at 50 °C. A sample volume of 1 mL was introduced into the GC-IMS system using the instrument sample loop. High-purity nitrogen (5.0, 99.999%) was used as both the GC carrier gas and the IMS drift gas. The drift gas flow was maintained at 100 mL/min throughout the analysis. The carrier gas flow was initially set to 5 mL/min and maintained for 15 s, followed by a linear increase to 30 mL/min at 120 s and subsequently to 150 mL/min at 180 s. The carrier gas flow was then maintained at 150 mL/min until the end of the 300 s analysis.

Following chromatographic separation, analytes were introduced into the IMS ionisation region; ionisation was achieved using a 370 MBq tritium source. The IMS was operated in positive ion mode and comprised a 10 cm drift tube maintained at 50 °C with an applied drift voltage of 5 kV. Ionised analytes were separated according to their mobility through the drift region against the counter-flowing nitrogen drift gas, resulting in characteristic drift times in addition to their chromatographic retention times.

GC-IMS data were acquired and processed using VOCal software. Each analysis generated a two-dimensional dataset comprising chromatographic retention time, ion mobility drift time, and signal intensity.

To assess sample validity and confirm that analysed samples were representative of exhaled breath, acetone and isoprene were used as endogenous breath markers, in addition to their role as an internal reference panel for the specificity analysis described below. For each measurement, both the peak volume and signal intensity of these compounds were evaluated using VOCal software [15]. A small number of individual samples could not be analysed successfully because of isolated data sampling failures, and were treated as missing: one oral sample (subject 007, *t*_1_) and two *t*_4_ samples (subjects 001 and 005), consistent with the sample sizes reported throughout the Results.

The primary analytes were propanal and 1-propanol, selected as candidate products of 1,2-PD metabolism. For each compound, the combined signal intensity of the corresponding monomer and dimer ion peaks was used for comparison (see **Figure 1**).

**Figure 1.**
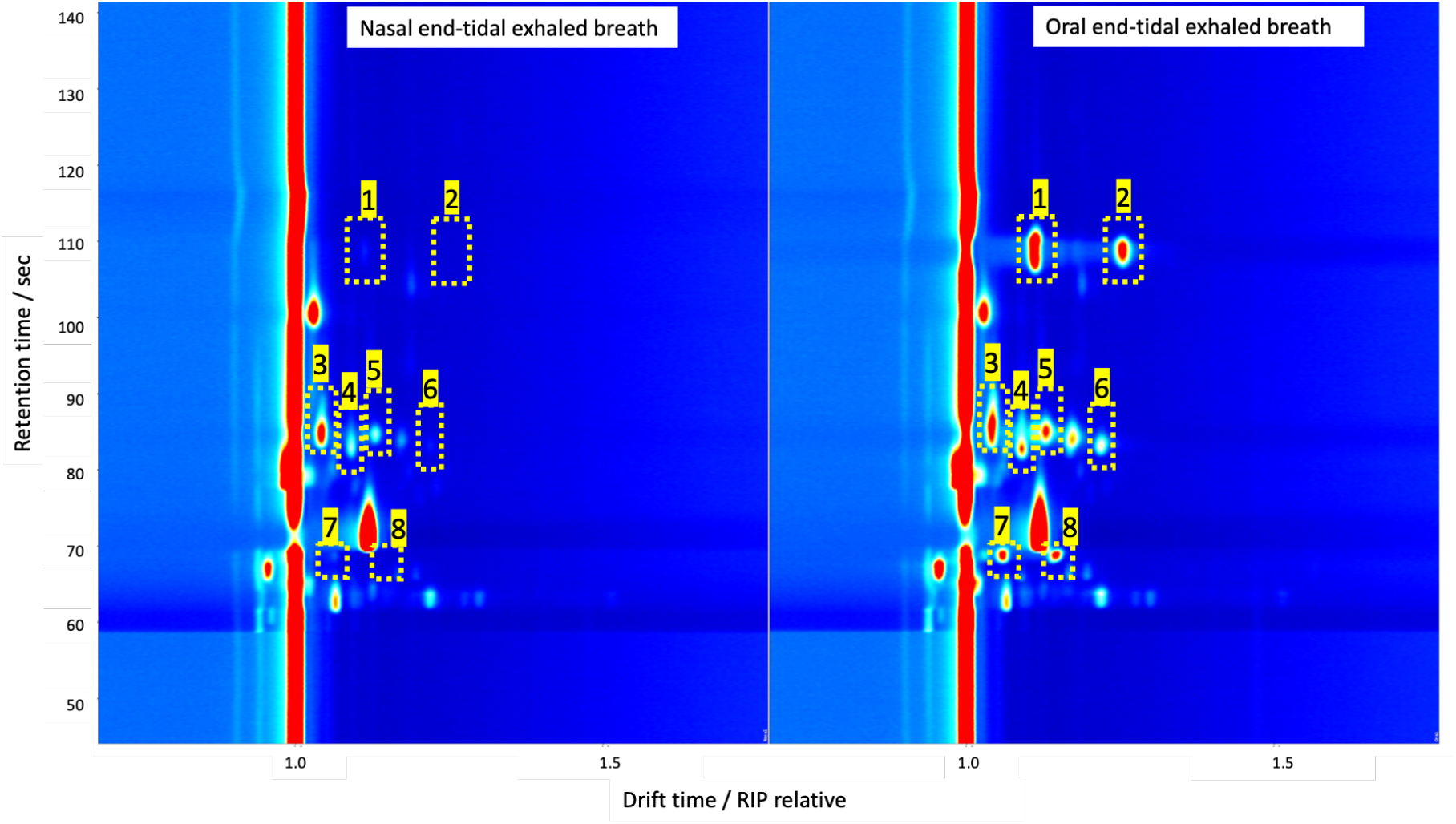
GC-IMS Chromatogram spectra of nasal end-tidal exhaled breath (left) and oral end-tidal exhaled breath (right). The x-axis provide the IMS drift time relative to a RIP located at 7.32 ms. The highlighted peaks are markedly more intense in the oral exhaled breath; these correspond to 1-propanol monomer and dimer (peaks 1 and 2), ethanol monomer and dimer (peaks 3 and 5), 2-propanol monomer and dimer (peaks 4 and 6), and propanal monomer and dimer (peaks 7 and 8).

Notably, acetol (hydroxyacetone) is a pathway-related analyte because it is represented in KEGG propanoate metabolism (map00640) and is associated with an alternative methylglyoxal-acetol route to 1,2-propanediol. It is not a principal downstream intermediate of the canonical B12-dependent *pdu* pathway from 1,2-PD to propanal and 1-propanol. Moreover, under the rapid GC-IMS method, the acetol signal was affected by interference from the aceticacid region; consequently, small or short-lived acetol changes may not have been resolved. Because its measurement was not sufficiently reliable, it was excluded from the formal statistical analysis. Isoprene, acetone, methanol, ethanol, and 2-propanol were monitored as reference VOCs to assess whether the intervention produced nonspecific changes in breath composition. These compounds were selected because previous route-resolved studies have shown that individual VOCs respond differently to changes in sampling route [6]. Dimethyl sulfide (DMS) was additionally included as a contrasting volatile sulphur compound: while hydrogen sulphide and methanethiol are commonly associated with intra-oral halitosis, DMS is the principal volatile implicated in extra-oral, blood-borne halitosis [16,17] and may therefore more closely reflect a systemic contribution to exhaled breath. Details of these compounds are provided in the Supplementary materials, in **Table S1**.

### 2.2 Data Analysis

This study was designed as a hypothesis-generating, mechanistic proof-of-concept investigation involving 10 participants and was not formally powered for clinical, diagnostic, or population-level inference. Effect sizes are therefore reported alongside adjusted *p*-values throughout and should be considered together with the magnitude, temporal pattern, and participant-level consistency of the observed responses. Because the cohort was small and each participant contributed repeated paired measurements across sampling routes and time points, non-parametric methods appropriate for matched longitudinal data were used. All analyses were performed in Python using the libraries SciPy, statsmodels, and scikit-learn [18].

For VOCs with resolvable monomer and dimer product-ion signals (1-propanol, propanal, ethanol, and 2-propanol), a combined volume under the peak surface was calculated as *I*_monomer_ + 2 *I*_dimer_. All inferential analyses were performed on log_10_(*x* + 1)-transformed intensities to reduce right-skewness and stabilise variance. Because this transformation is strictly monotonic for non-negative intensity values, it preserves the ordering and direction of paired differences. Ratios and fold changes are reported in the original, non-transformed peak volume intensity scale to facilitate interpretation.

Principal component analysis (PCA) was adopted for aiding the multivariate analysis. It used log10(x+1)-transformed, standardised (unit-variance-scaled) intensities throughout. Conventional null-hypothesis significance testing (H_0_: no difference) can flag a compound as statistically significant on the strength of a highly consistent direction across volunteers alone, even when the underlying magnitude of change is biologically trivial. To distinguish compounds substantially, rather than merely detectably, affected by the intervention, three complementary minimum-effect procedures were applied, following the general logic of equivalence testing (shifting the tested null value away from zero to a pre-specified, biologically motivated threshold, rather than testing against no difference).

Changes relative to the pre-intervention baseline were assessed for the primary analytes using repeated-measures comparisons, with post hoc comparisons against *t*_pre_ where appropriate. Specifically, each volunteer’s log_2_(oral/nasal) ratio at *t*_pre_ was shifted by log_2_(3/2) and tested with a one-sided Wilcoxon signed-rank test, testing whether oral levels were reliably at least 1.5-fold higher than nasal, Holm–Bonferroni corrected across the eight compounds at this time point.

Within-route temporal effects were assessed with the non-parametric Friedman test across *t*_pre_-t_3_, with Kendall’s W as the primary effect-size measure (0 = no association between time and rank, 1 = perfect association). Holm-Bonferroni correction was applied across all 16 tests (8 compounds × 2 routes). A compound×route combination was classified as substantially time-dependent only if the 95% bootstrap confidence interval for Kendall’s W (5000 resamples, resampling volunteers with replacement) lay entirely above W = 0.5, a conventional threshold for a moderate-to-strong effect.

A complementary Baseline→Peak→End summary (Peak fixed at t0, End fixed at t3) for all compounds was implemented to quantify the temporal changes, with post-hoc paired one-sided Wilcoxon signed-rank tests, Holm–Bonferroni-corrected within each compound’s three-comparison family, and effect sizes reported as matched-pairs rank-biserial correlation coefficients (r; −1 = perfect decrease, +1 = perfect increase across all volunteers). One-sided Wilcoxon signed-rank test was applied to each volunteer’s log_2_(peak/baseline) ratio, shifted by log_2_(1.10), testing H_0_: median rise ≤ 10% against H_1_: median rise > 10%, Holm-corrected across the eight compounds. This criterion was pre-specified as a round, biologically conservative threshold (a 10% rise) rather than tuned to the data;

## 3. Results

Before evaluating the response to the 1,2-PD intervention, the reproducibility of the baseline oral-nasal route effect reported in our previous study [6] was assessed in this independent cohort. This analysis served two purposes: first, to confirm that the paired oral and nasal sampling protocol reproduced the expected route-dependent pattern before intervention-specific effects were considered; and second, to establish a pre-intervention reference for interpreting the subsequent temporal analyses. Oral and nasal end-tidal breath samples collected at *t*_-1_ were compared for all eight monitored VOCs. Rather than testing whether nasal/oral differed from 1 (which even a small but consistent shift can satisfy), we tested whether it was reliably below 2/3, i.e. VOC level in the oral sample at least 1.5-fold higher than in the nasal exhaled breath sample. This was done with a one-sided Wilcoxon signed-rank test (as detailed in the Methods), with Holm-Bonferroni correction across the eight compound-specific comparisons (see **Figure 2**; **Table 1**, “Pre” column).

**Table 1.** Nasal/oral ratio (%) by compound and time point, and minimum-effect test: significance of “ratio < 2/3 (66.7%)” (one-sided Wilcoxon signed-rank test on log2[(nasal+1)/(oral+1)], shifted by log2(2/3); Holm–Bonferroni corrected within each time point, 8 compounds). n = 10 at pre, t0, t2 and t3; n = 9 at t1; n = 8 at t4. * p < 0.05, ** p < 0.01 (Holm-adjusted).

| **VOC** | *t*<sub>pre</sub> |  | *t*<sub>0</sub> |  | *t*<sub>1</sub> |  | *t*<sub>2</sub> |  | *t*<sub>3</sub> |  | *t*<sub>4</sub> |  |
|---|---|---|---|---|---|---|---|---|---|---|---|---|
|  | **Ratio** | **Sig.** | **Ratio** | **Sig.** | **Ratio** | **Sig.** | **Ratio** | **Sig.** | **Ratio** | **Sig.** | **Ratio** | **Sig.** |
| **1-Propanol** | 4% | ** | 3% | ** | 4% | * | 3% | ** | 4% | ** | 2% | * |
| **Propanal** | 43% | n.s. | 9% | ** | 12% | * | 45% | n.s. | 46% | n.s. | 51% | n.s. |
| **Ethanol** | 33% | ** | 40% | * | 34% | * | 38% | ** | 38% | ** | 43% | * |
| **2-Propanol** | 43% | * | 57% | * | 44% | * | 43% | * | 43% | n.s. | 48% | n.s. |
| **DMS** | 66% | n.s. | 68% | n.s. | 77% | n.s. | 70% | n.s. | 64% | n.s. | 56% | n.s. |
| **Acetone** | 95% | n.s. | 87% | n.s. | 91% | n.s. | 91% | n.s. | 93% | n.s. | 93% | n.s. |
| **Isoprene** | 75% | n.s. | 83% | n.s. | 89% | n.s. | 95% | n.s. | 84% | n.s. | 67% | n.s. |
| **Methanol** | 104% | n.s. | 101% | n.s. | 106% | n.s. | 105% | n.s. | 102% | n.s. | 105% | n.s. |

**Figure 2.**
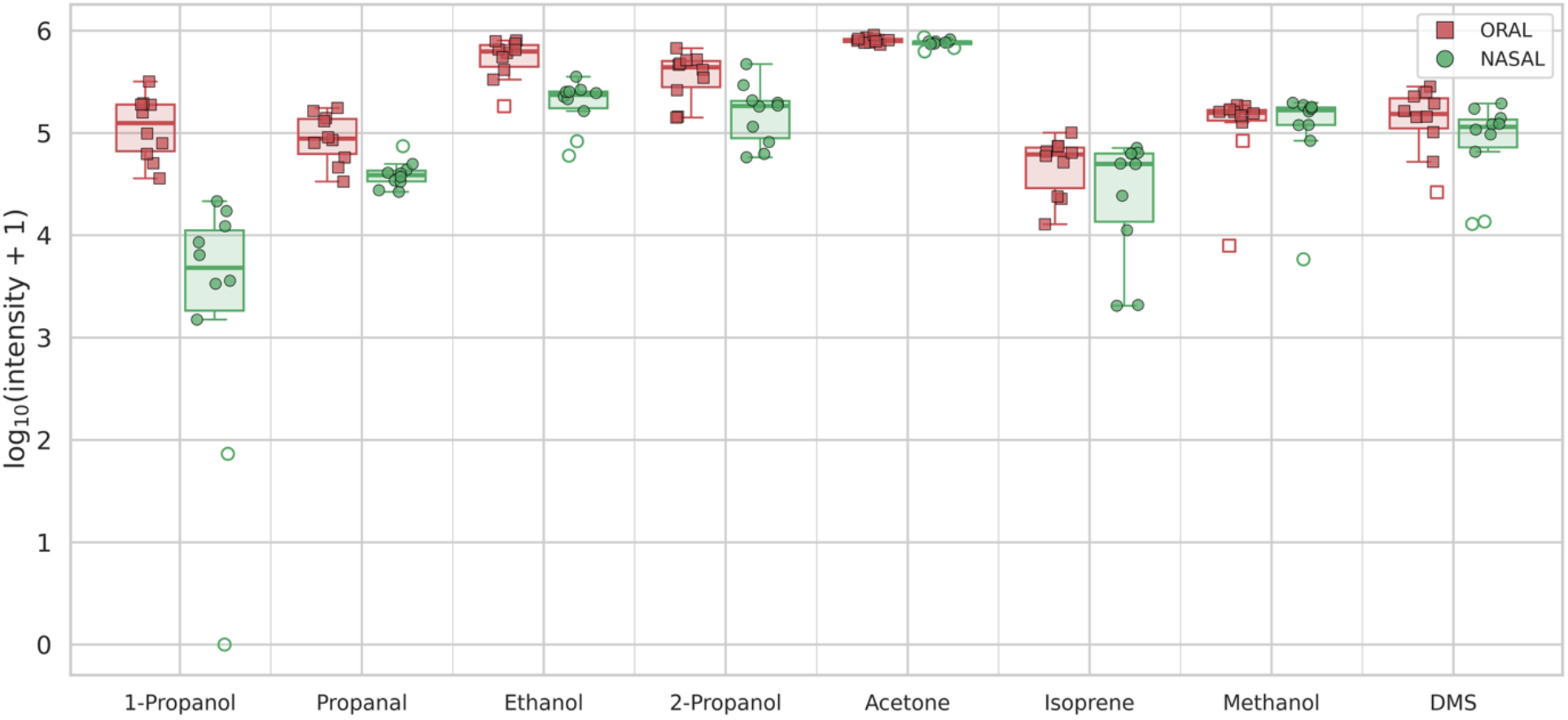
Boxplots of the pre-intervention dataset. The results are in agreement with former study [6].

Oral administration of 1,2-PD produced a rapid increase in orally exhaled propanal and 1-propanol signals. For both compounds, the largest oral responses occurred immediately after the intervention at *t*_0_ and remained elevated at *t*_1_. Signals subsequently declined during *t*_2_-*t*_4_ towards the pre-intervention state (see **Figure 4 and S2**.).

To explore the multivariate structure of the dataset, and to identify which VOCs were most closely associated with sampling route and with the time-course of the intervention, principal component analysis (PCA) was applied to the panel of monitored VOCs, log-transformed and standardised throughout. **Figure 3** panel a) shows the global ordination (all sample types pooled) whilst panel b) shows the results when only the two main intervention-associated VOCs and the two principal systemic VOCs.

**Figure 3.**
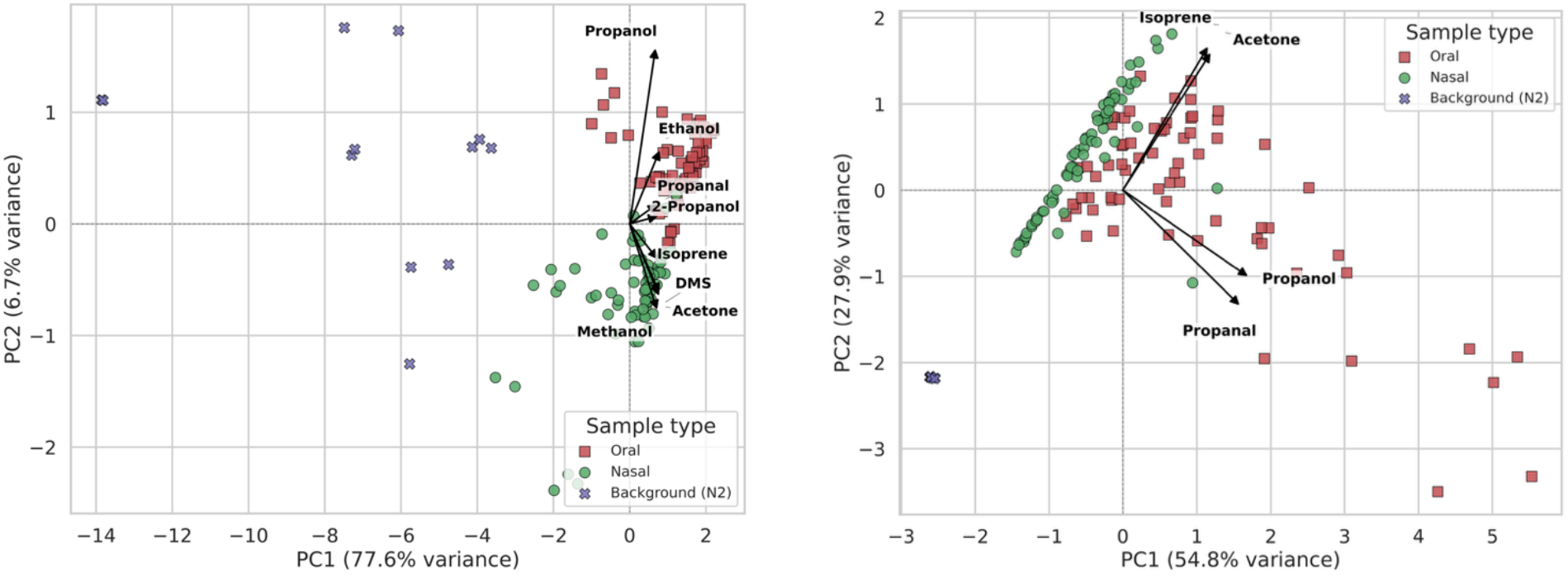
(a) PCA biplot for all the pooled time-points. Aceton and isoprene are aligned parallel to the separation of the nasal and oral clusters of points, as expected from systemic compounds. (b) PCA bi-plot plot selecting only four out of the eight monitored VOCs; namely, propanol and 1-propanol (intervention-associated) and acetone, and isoprene (systemic compounds, unaffected by the intervention).

**Figure 4.**
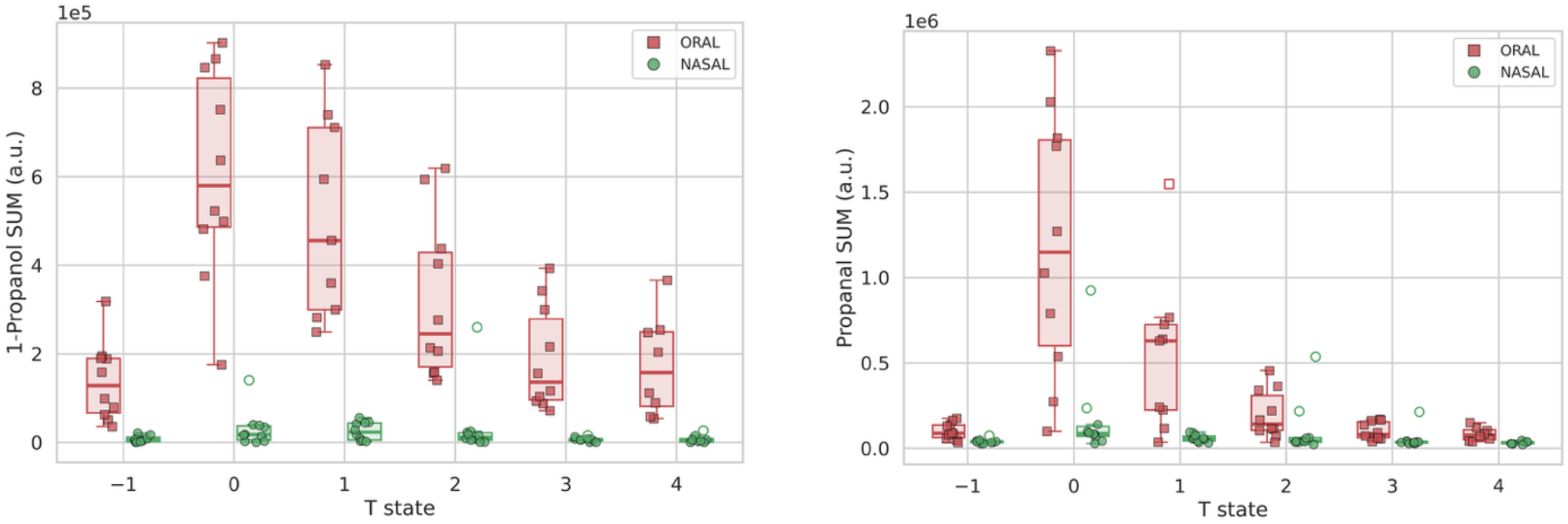
Boxplots showing the route-resolved variation of the intensities of 1-propoanol and propanal for the participants across the protocol time-points. T states on the x-axis denote: -1, pre-exposure baseline (*t*_pre_) collected immediately before the rinse; 0, immediately after exposure (*t*_0_); and 1–4, approximately 15, 30, 45 and 60 minutes after exposure (*t*_1_-*t*_4_).

Across these representations, 1-propanol and propanal consistently emerged as the VOCs most strongly associated with both the oral/nasal separation and the post-intervention trajectory, whereas acetone and isoprene tracked an axis shared along the background/breath samples separation (**Figure 3b**).

Nasal propanal and 1-propanol signals remained comparatively stable, producing a clear early oral-nasal signal intensity separation that diminished during washout as oral signals returned towards baseline. The response was therefore characterised by an early oral maximum followed by progressive washout. This temporal and exhalation route-specific pattern is consistent with a local response to oral 1,2-PD exposure rather than an immediate systemic change expected to affect both sampling routes.

To ultimately determine which compounds were substantially, affected by the intervention, we applied a minimum-effect criterion to the time-course analysis: a compound×route combination was classified as substantially time-dependent only if the 95% bootstrap confidence interval for Kendall’s W lay entirely above the W = 0.5 threshold (**Figure 5**). Of the sixteen compound×route combinations tested, only oral 1-propanol (W = 0.86, 95% CI 0.78-0.97) and oral propanal (W = 0.80, 95% CI 0.67-0.97) met this criterion. Oral acetone (W = 0.46, 95% CI 0.26-0.82) and nasal 1-propanol (W = 0.47, 95% CI 0.25-0.79) were both nominally significant under the conventional Friedman test (*p*_Holm_ < 0.05) but had confidence intervals spanning the threshold, and were therefore not classified as substantially affected; this distinguishes a small but statistically detectable shift, achievable given a sufficiently consistent direction across ten volunteers, from a change large enough to be confirmed as such independent of the specific significance threshold chosen. Notably, nasal 1-propanol’s own significant time effect indicates that 1-propanol also rises modestly in nasal breath following the intervention; because both routes move together, the oral/nasal ratio for 1-propanol remains essentially constant across time points (**Table 1**) even though its absolute oral concentration is clearly time-dependent. Propanal shows the converse pattern: its nasal signal is not significantly time-dependent (W = 0.30, *p*_Holm_ = 0.22), so its much stronger oral response produces a pronounced, transient collapse in the oral/nasal ratio specifically around the time of the intervention (**Table 1**) rather than a constant offset. Acetone, despite its nominal significance, never reached even half the effect size of either primary analyte, and its rise from baseline (increment ∼ 6% of baseline; **Table 2**) was modest, further supporting its exclusion from the set of compounds substantially affected by the intervention.

**Table 2.** Temporal changes (expressed in percentage increment or reduction) in end-tidal oral exhaled breath VOCs. n.s. = not significant (p > 0.05, Holm-adjusted within each VOC’s 3-comparison family), * p < 0.05, ** p < 0.01. r = matched-pairs rank-biserial correlation. † Baseline → Peak rise additionally confirmed as reliably exceeding a 10% minimum-effect threshold (one-sided Wilcoxon signed-rank test on the shifted log-ratio, Holm-corrected across the eight compounds; p_*Holm*_ *= 0*.*0078)*.

| VOC | **Baseline (t<sub>pre</sub>) → Peak (t<sub>0</sub>)** | **Peak (t<sub>0</sub>) → End (t<sub>3</sub>)** | **Baseline (t<sub>pre</sub>) → End (t<sub>3</sub>)** |
|---|---|---|---|
| **1-Propanol** | +381.8%, ↑** (r=1.00), † | -70.3%, ↓** (r=-1.00) | +76.7%, ↑* (r=0.71) |
| **Propanal** | +1413.4%, ↑** (r=1.00), † | -91.5%, ↓** (r=-1.00) | +18.0% (n.s.) |
| **Acetone** | +5.8%* (r=0.85) | -4.9%, ↓* (r=-0.96) | +1.1% (n.s.) |
| **Isoprene** | +5.0% (n.s.) | -16.1% (n.s.) | -2.3% (n.s.) |
| **Methanol** | -2.4% (n.s.) | +0.9% (n.s.) | +2.8% (n.s.) |
| **DMS** | -9.4% (n.s.) | +16.2% (n.s.) | +3.3% (n.s.) |
| **Ethanol** | -10.9% (n.s.) | -1.4% (n.s.) | -12.9% (n.s.) |
| **2-Propanol** | -11.0% (n.s.) | +8.9% (n.s.) | -13.9% (n.s.) |

**Table 3.** Overview of the different roles and mechanisms, with evidence linking salivary glycans, oral microbial metabolism and breath volatiles.

| Evidence domain | Representative organism or system | What the study demonstrates | What it does not demonstrate for this study | Key reference |
|---|---|---|---|---|
| **Fucosylated salivary host substrates** | Human salivary MUC5B and MUC7 | Fucosylated O-glycan structures occur on both salivary mucins | Release of fucose by oral bacteria or conversion to 1,2-PD *in vivo* | [22,23] |
| **Oral glycoprotein-sugar metabolism** | Human saliva plus plaque/salivary sediment | Oral bacteria can degrade salivary glycoprotein sugars, including fucose | Identity of the responsible taxa, free-fucose kinetics, and downstream metabolic products in contemporary oral biofilms | [13]11/6/26 2:45:00 PM |
| **Fucose to 1,2-PD formation** | *Roseburia inulinivorans* | Fucose-associated expression of fucose and propanediol pathways, with propanol and propionate production | Operation of this pathway in oral communities | [24] |
| **1,2-PD utilisation** | *Propionibacterium freudenreichii* | Anaerobic 1,2-PD utilisation yielding propionate and 1-propanol | Relevance to oral biofilms | [25,26] |
| **Oral 1,2-PD-use capacity** | *Streptococcus sanguinis* genome | A *pdu* cluster and B12 biosynthesis genes are present | In vivo expression, substrate use, or breath VOC production | [27] |
| **Human oral response** | Orally exhaled air gets enriched with local production of VOCs. | Oral 1,2-PD exposure produces route-localised propanal and 1-propanol signals | Microbial identity, enzyme activity, endogenous 1,2-PD source, and mucin-carbon flux | Present study |

**Figure 5.**
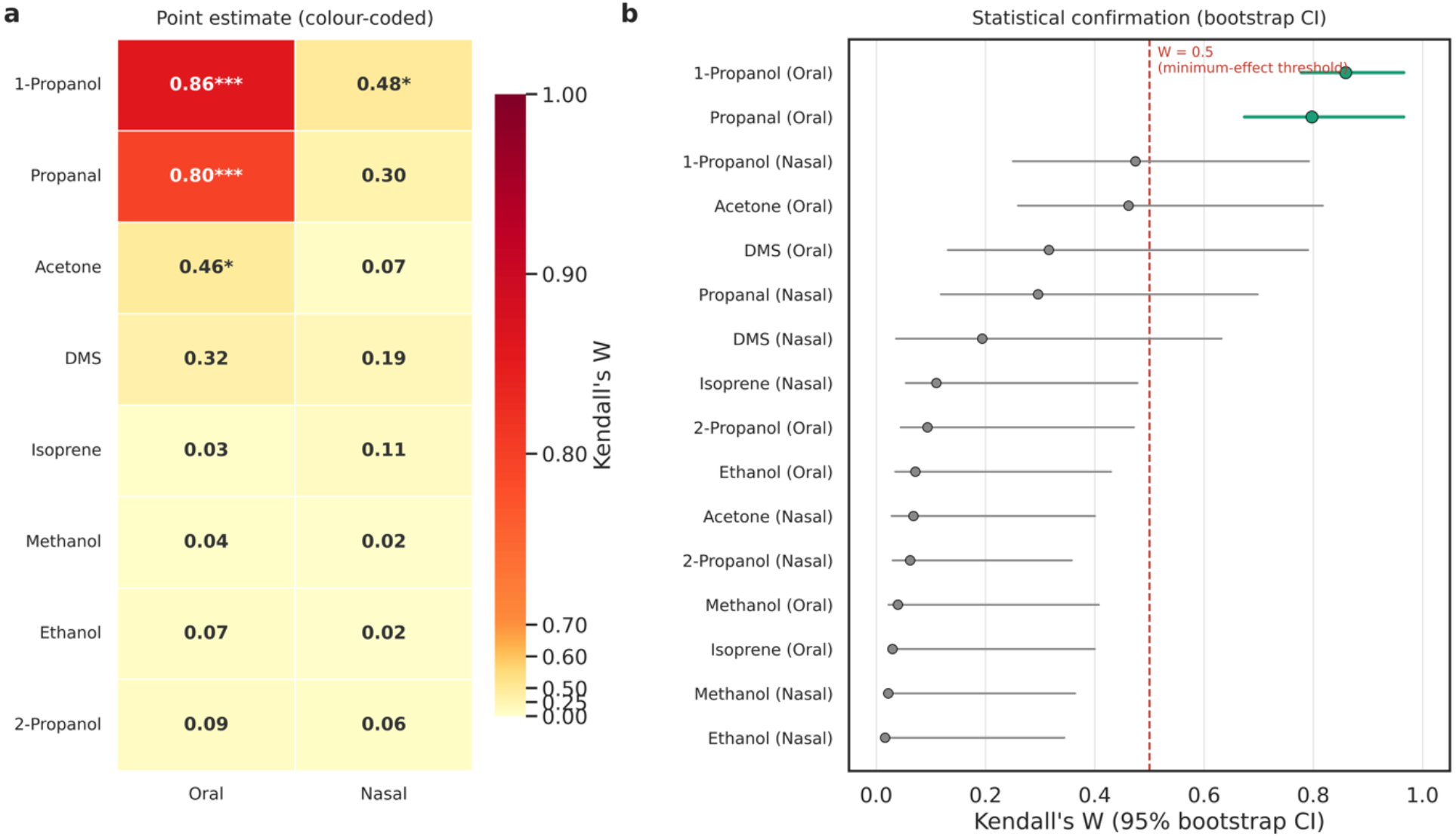
Time-course effect size by compound and route (Friedman test, t_pre_-t_3_). (a) Kendall’s W point estimates, colour-coded on a scale expanded above W = 0.5 (large effect threshold) to emphasise the two compounds exceeding the minnimum-effect threshold; asterisks denote significance after Holm-Bonferroni correction across all sixteen tests (* p < 0.05, ** p < 0.01, *** p < 0.001). (b) The same effect sizes with 95% bootstrap confidence intervals (5000 resamples, resampling volunteers); green markers denote combinations whose interval lies entirely above the W = 0.5 threshold (red dashed line).

**Table 2** summarises the temporal response of all eight VOCs in oral exhaled breath using a complementary approach, with *Peak* fixed at *t*_0_ and *End* fixed at *t*_3_ for every compound. Specifically, *t*_3_ was selected to retain complete paired data for all participants (*n* = 10); use of *t*_4_ would have resulted in reduced and non-uniform sample sizes because of missing observations. 1-propanol and propanal were the only two compounds showing a rise from baseline to peak that was both large in magnitude and present in every single volunteer (1-propanol: median +382% increase, r = 1.00; propanal: +1413% increase, r = 1.00; both *p*_Holm_ < 0.01), followed by a significant decline from peak to *t*_3_ (both r = −1.00). Conversely, acetone showed a much smaller, less uniform variation (median +5.8% increase, r = 0.85); the remaining five compounds showed no significant change at any comparison. This distinction is consistent with the proposed biochemical pathway: acetone is not a reported intermediate or product of 1,2-PD degradation via the coenzyme B12-dependent propanediol-utilisation route (KEGG map00640). Its modest, individually inconsistent rise is more plausibly attributable to a small systemic (e.g., ventilation/perfusion) perturbation than to direct metabolic processing of the administered substrate.

The overall direction of the response was consistent, but its magnitude varied between participants. The greatest between-participant variation occurred at t_0_ and t_1_, when the administered substrate was most recently present in the oral compartment. Participants showed pronounced increases in both end-tidal oral exhaled propanal and 1-propanol, with some showing a more moderate response than others. Nasal signals displayed little-to-no observable temporal variation across the same sampling sequence. Differences in breath flow, salivary flow, oral retention, biofilm structure, substrate access, microbial community composition, and local physicochemical conditions could contribute to this variation. These factors were not measured concurrently and therefore remain possible explanations rather than demonstrated determinants of the response.

Acetone and isoprene did not show a consistent post-intervention increase or decrease in either sampling route. Their relative stability contrasted with the transient oral responses of propanal and 1-propanol, suggesting that the intervention did not produce a broad change in exhaled VOC abundance.

## 4. Discussion

The intervention produced a selective and transient increase in oral propanal and 1-propanol, with little corresponding nasal response. This compartment-specific pattern supports the interpretation that 1,2-PD was acted upon within the oral environment or that its immediate volatile products accumulated there. The temporal profile is compatible with rapid access to an orally administered substrate, followed by salivary clearance, and loss of volatile products during the observation time window.

The results extend previous route-resolved observations [6]. Oral and nasal sampling can produce different VOC profiles [8,9], and oral microorganisms can transform substrates within the mouth and contribute volatile compounds to mouth-exhaled breath [4,5]. Previous work also indicated oral microbiome activity as an important consideration when interpreting breath VOCs [6]. The present intervention adds a targeted substrate perturbation: direct oral exposure to 1,2-PD was followed by an increase in 1-propanol together with propanal.

The stable nasal measurements argue against a large systemic contribution within the short observation window. Furthermore, the stability of the other VOCs (i.e., isoprene, acetone, methanol, DMS, ethanol and 2-propanol) provides an additional internal comparison. The intervention did not produce a generalised change in breath VOC abundance but instead produced a selective response involving compounds associated with the administered substrate. As discussed in the previous sections, the acetol channel was limited by chromatographic interference.

### Oral Microbial Ecology and Candidate Endogenous Substrates

The oral cavity contains multiple microbial habitats, including saliva, the tongue, mucosal surfaces, and dental biofilms [19,20]. These habitats support interacting microbial communities whose composition and metabolic activity can vary across individuals and ecological niches [21].

Salivary mucins are heavily O-glycosylated glycoproteins that contribute to the physicochemical properties and protective functions of the oral mucosal environment and can influence oral microbial community composition and function (Tabak, 1995; Derrien et al., 2010, Wu 2023). Specifically, work has further shown that oral commensals can utilise salivary MUC5B as a sole nutrient source and exhibit cooperative metabolism during early biofilm formation (Robertsson et al., 2023); fucosylated O-glycan structures have also been characterised on human salivary MUC7 (Chaudhury et al., 2016). Mucin glycan are therefore plausible host-derived contributors to the fermentable substrate pool.

These findings support the general ecological premise that host-derived substrates can contribute to oral microbial metabolism. They do not, however, demonstrate that mucin-derived fucose is converted to 1,2-PD in the human oral cavity or that such a pathway produces the baseline 1-propanol signal observed in fasting participants. The proposed sequence (**Figure 6**) should therefore be treated as a hypothesis:

host-derived oral substrates → candidate 1,2-PD formation → propanal → 1-propanol.

**Figure 6.**
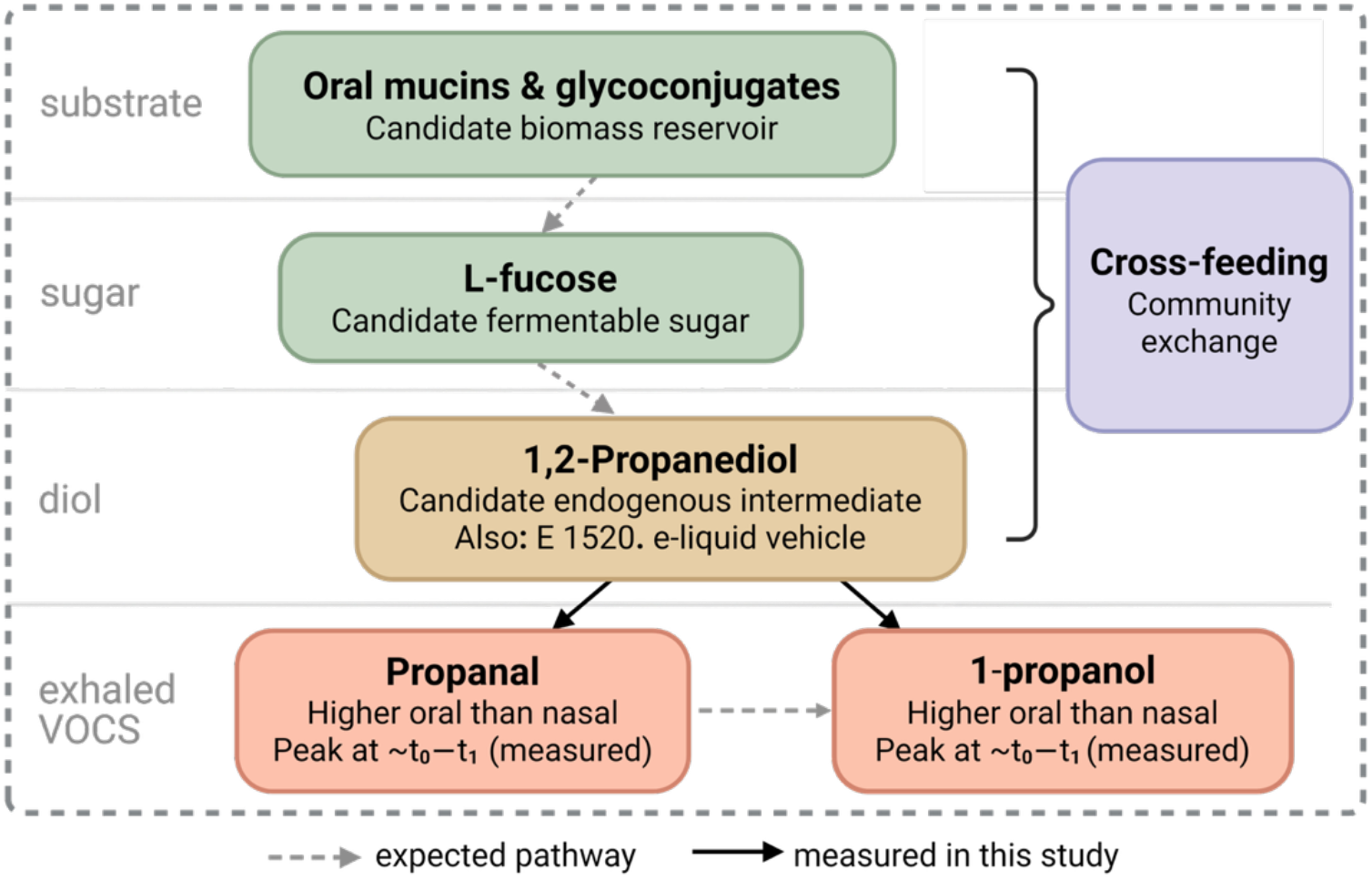
Diagram depicting the proposed framework of the mechanistic biochemical pathways involved. The appearance of propanal and 1-propanol following oral 1,2-PD exposure is supported by the present intervention (solid arrows), whereas the upstream steps linking salivary mucin glycans through fucose to 1,2-PD are inferred from bacterial systems characterised in other niches (dashed arrows).

The present intervention directly tested only the downstream response to exogenous 1,2-PD. It supports the possibility that the oral environment can transform or otherwise process 1,2-PD into volatile products, but it does not establish the endogenous origin of 1,2-PD.

A distributed pathway involving several microbial taxa is also plausible. Mucin degradation, sugar metabolism, and downstream propanediol utilisation need not occur within a single organism. Oral biofilms contain interacting microbial networks and spatially distinct microenvironments [28]. Community-level metabolite production can indeed reflect cooperation or cross-feeding between organisms rather than the activity of one isolated species [29]. In principle, 1,2-PD may also arise from a hepatic or systemic source independent of salivary mucin-derived fucose: acetone can be converted to acetol and subsequently to 1,2-PD [30]. However, there is currently no direct evidence that acetone-derived 1,2-PD reaches saliva at concentrations sufficient to support local microbial propanediol utilisation and baseline oral 1-propanol production.

### Implications for Breath-Biomarker Research

The findings have direct implications for breath-biomarker interpretation. A volatile compound detected in exhaled breath may contain contributions from systemic circulation, airway tissues, and oral microbial activity [3,31].

Fasting reduces recent dietary exposure but certainly does not eliminate oral microbial activity. Host-derived substrates remain present in the oral environment, and mucins can interact with and support oral microbial communities [34,35]. Fasting should therefore be treated as a control for recent food intake, not as proof that all oral microbial volatile contributions have been removed.

Notably, 1,2-Propanediol (1,2-PD) is detectable in human saliva, with reported basal concentrations of approximately 0.4 mg/L in non-users of tobacco or nicotine products [32] and approximately 2.2 mg/L in the control group of a salivary metabolomics study [33]. Combined with the observed fasting-persistent oral enrichment of 1-propanol [6] and the predominantly oral-specific conversion of exogenous 1,2-PD to propanal and 1-propanol shown here, this supports the interpretation that baseline oral 1-propanol arises, at least in part, from microbial conversion of salivary 1,2-PD. Salivary 1,2-PD may therefore provide a continuous, low-level substrate source sustaining baseline oral 1-propanol production.

In addition to endogenous and dietary sources, 1,2-propanediol is widely encountered as propane-1,2-diol, a permitted food additive and carrier (E 1520), and as a major constituent of many electronic-cigarette liquids and tobacco products [32]. Hence, these exposures should be considered as potential pre-analytical confounders in exhaled breath studies.

Matched nasal and oral sampling provides one practical means of identifying this contribution. In the present study, nasal end-tidal measurements acted as a comparator for the oral response. More generally, end-tidal nasal breath sampling may help reduce direct exposure to locally produced oral volatiles when the scientific objective is to investigate predominantly systemic or pulmonary VOCs.

Oral cleansing may also be used experimentally to assess oral contributions. However, cleansing procedures require careful standardisation because they can change the oral environment and alter VOC measurements independently of systemic physiology [36]. Oral contamination and pre-analytical variation should therefore be considered when putative breath biomarkers are validated [5,6,37].

The same biology may eventually be exploited rather than merely controlled. Substrate-challenge breath tests designed to probe oral microbial metabolic capacity could potentially provide functional information relevant to oral health. This possibility remains speculative and requires validation against clinical measures, microbiome composition, and direct metabolite measurements.

### Analytical Limitations and Future Directions

Several limitations constrain interpretation. The cohort was small, the observation period was brief, and the experiment did not include an independent control intervention. The oral microbiome, salivary flow and breath flow, pH, mucin concentration, and oral-health status were not measured concurrently. Consequently, the biological sources of inter-individual variation remain unresolved.

The study also tested exogenous 1,2-PD and therefore cannot establish that endogenous host-derived substrates sustain baseline oral 1-propanol production. Although the complete pathway has not yet been demonstrated in vivo, the present findings complete a convergent body of evidence in which each proposed link is supported independently: fucosylated salivary mucins provide a plausible substrate architecture, oral communities can process salivary glycoprotein sugars, fucose and 1,2-propanediol occur in saliva, and exogenous 1,2-propanediol elicits route-localised propanal and 1-propanol responses.

To our knowledge, the complete trophic pathway proposed here (see **Figure 6**), comprising microbial liberation of fucose from salivary mucins, fermentation of fucose to 1,2-propanediol, and subsequent utilisation of 1,2-propanediol to form propanal and 1-propanol, has not been directly demonstrated in an oral microbial community. Individual components of this pathway are nevertheless supported by previous observations, including bacterial degradation of salivary glycoprotein sugars, the occurrence of 1,2-propanediol-utilisation loci in oral-associated bacteria, and evidence that oral microorganisms contribute to 1-propanol in saliva and exahled breath. An analogous mucin-derived 1,2-propanediol cross-feeding cascade has been experimentally demonstrated in the intestinal microbiota, making its occurrence in the oral ecosystem biologically plausible but currently unverified.

The proposed oral pathway could be investigated *in vitro* by incubating saliva-derived or dental-biofilm communities in parallel with characterised salivary MUC5B, L-fucose or 1,2-PD under controlled conditions. Time-resolved measurements of free fucose, 1,2-PD and propionate in the liquid phase, together with headspace propanal and 1-propanol, would help assess the successive stages of mucin utilisation, fucose fermentation and downstream 1,2-PD metabolism.

Future work should also determine whether exhalation route-dependent propanal and 1-propanol responses are associated with oral health, dysbiosis, or periodontal inflammation. Reviews of oral microbiota and metabolites describe relationships between community changes, microbial metabolites, and oral health or disease [38]. However, such associations should not be assumed for the present compounds without direct clinical validation.

## 5. Summary and conclusions

Controlled oral exposure to 1,2-PD produced a rapid increase in propanal and 1-propanol in oral, but not nasal, end-tidal exhaled breath. The early maximum, subsequent washout of these two volatiles, and relative stability of acetone and isoprene support a local, substrate-associated response. These findings strengthen the interpretation that oral microbial activity contributes to route-dependent breath VOCs. This study completes the central experimental arc initiated by the original exhalation route-resolved observation. Firstly we identified a subset of VOCs (including 1-propanol, ethanol, and acetoin) that differed markedly between oral and nasal end-tidal breath [6]. Then we showed that sub-gram oral glucose exposure elicited rapid acetoin and ethanol responses, supporting a local fermentation-associated origin for these VOCs [4], while leaving the route-dependent 1-propanol signal unresolved. The present 1,2-PD challenge addresses this gap by demonstrating a transient, oral-specific response of propanal and 1-propanol to a pathway-informed candidate substrate.

The result should be interpreted as evidence for a local oral response to exogenous 1,2-PD, while leaving the endogenous source of 1,2-PD and the responsible microbial taxa for future direct investigation. The proposed pathway linking salivary mucin-derived fucose to 1,2-propanediol, propanal, and 1-propanol remains to be demonstrated directly. Nevertheless, salivary mucins represent a plausible continuously available host-derived biomass reservoir for oral microbial metabolism, because dental-plaque communities can degrade and utilise salivary proteins (e.g., MUC5B and 7) in the absence of recent dietary carbohydrate exposure. Fasting should therefore not be assumed to eliminate oral microbial contributions to exhaled VOC profiles. Nasal end-tidal sampling provides a practical comparator when the primary objective is to characterise systemic VOCs, specifically in untargeted biomarker discovery studies [3].

*In vivo* incubations of oral communities with mucin glycoproteins, L-fucose or 1,2-PD substrates is the clearest next step to test whether this mucin-associated metabolic sequence, described primarily in gut communities, operates in the oral cavity and contributes to the production of exhaled propanal and 1-propanol.

More broadly, route-dependent oral VOCs may eventually be useful as functional indicators of microbial activity rather than being treated solely as sources of biomarker confounding. Before such applications are possible, their biochemical origin, reproducibility, biological specificity, and clinical relevance must be established through controlled longitudinal studies integrating exhalation route-resolved breath analysis with saliva metabolomics and oral microbial profiling.

## Acknowledgments

We wish to thank Univ.-Prof. Dr. Chris A. Mayhew, Director of the Institute for Breath Research, University of Innsbruck, Austria, for his strong support of this research project through his scientific input and the provision of instrumental and consumable resources. We also gratefully acknowledge his valuable contributions to the initial projects that ultimately led to this manuscript. Support for Anesu Chawaguta’s doctoral research was provided through the Austrian Research Promotion Agency (FFG) program IKT der Zukunft, IKT der Zukunft-Resilienz und Distancing (DEVICE). Funding for Dr. Lorenzo Petralia was provided by the EU Horizon Innovation Action ONELAB (Grant Agreement No. 101073924; HORIZON-CL3-2022-DRS-01-05).

## Data Availability Statement

The datasets generated and/or analysed during the current study are available from the corresponding author upon reasonable request.

## Conflict of Interest

Dr Daniel Sanders is a Research and Development Scientist at G.A.S. Gesellschaft für analytische Sensorsysteme mbH (Dortmund, Germany), which develops, manufactures, and commercializes the GC-IMS instrument utilised in this study. All remaining authors declare no competing interests.

## Compliance with ethical standards

Ethical approval for this study was obtained from the Institutional Ethics Committee of the Medical University of Innsbruck (EK Nr.: 1181/2017). Prior to sample collection, all participants received information regarding the purpose and procedures of the study and provided written informed consent. The research was conducted in accordance with the ethical principles outlined in the 1975 Declaration of Helsinki.

## Notes

### Competing Interest Statement

Dr Daniel Sanders is a Research and Development Scientist at G.A.S. Gesellschaft fuer analytische Sensorsysteme mbH (Dortmund, Germany), which develops, manufactures, and commercializes the GC-IMS instrument utilised in this study. All remaining authors declare no competing interests.

