## Supplementary Material for "Interventional breath analysis reveals oral microbial conversion of 1,2-Propanediol into Propanal and 1-Propanol"

Compound identities were assigned from gas chromatographic retention time and ion mobility drift time, expressed as the derived reduced ion mobility, $K_0$ ($cm^2\ V^{-1}\ s^{-1}$), under the GC-IMS operating conditions described in Methods. Four of the eight monitored VOCs (1-propanol, propanal, ethanol and 2-propanol) resolved into two distinct product-ion clusters, corresponding to the monomer (M) and proton-bound dimer (D) species commonly observed by IMS for small, polar volatiles at sufficient concentration; both were quantified and combined (monomer + 2×dimer) for statistical analysis, as stated in the main text. The remaining four VOCs (acetone, isoprene, methanol, and DMS) were each observed as a single product-ion channel. Retention time, drift time and $K_0$ for all twelve product-ion channels are summarised in **Table S1** below.

| Compound | Formula | Retention Time (s) | Drift Time (ms) | $K_0$ ($cm^2V^{-1}s^{-1}$) |
|---|---|---|---|---|
| **1-Propanol M** | $C_3H_8O$ | 110.84 | 8.110 | 1.87 |
| **1-Propanol D** | $C_3H_8O$ | 110.84 | 9.162 | 1.65 |
| **Propanal M** | $C_3H_6O$ | 69.59 | 7.729 | 1.96 |
| **Propanal D** | $C_3H_6O$ | 69.59 | 8.368 | 1.81 |
| **Ethanol M** | $C_2H_6O$ | 86.64 | 7.624 | 1.99 |
| **Ethanol D** | $C_2H_6O$ | 86.64 | 8.267 | 1.83 |
| **2-Propanol M** | $C_3H_8O$ | 84.37 | 7.963 | 1.90 |
| **2-Propanol D** | $C_3H_8O$ | 84.37 | 8.914 | 1.70 |
| **Acetone** | $C_3H_6O$ | 72.88 | 8.175 | 1.85 |
| **Isoprene** | $C_5H_8$ | 64.27 | 8.919 | 1.70 |
| **Methanol** | $CH_4O$ | 81.15 | 7.713 | 1.97 |
| **DMS** | $C_2H_6S$ | 67.76 | 6.985 | 2.17 |

**Table S1.** Compound identification table reporting GC-IMS retention time (in seconds), product-ion drift time (in ms), and reduced ion mobility $K_0$ (in $cm^2V^{-1}s^{-1}$) for the eight monitored VOCs. Compounds with resolvable monomer (M) and proton-bound dimer (D) product ions are listed as separate rows.

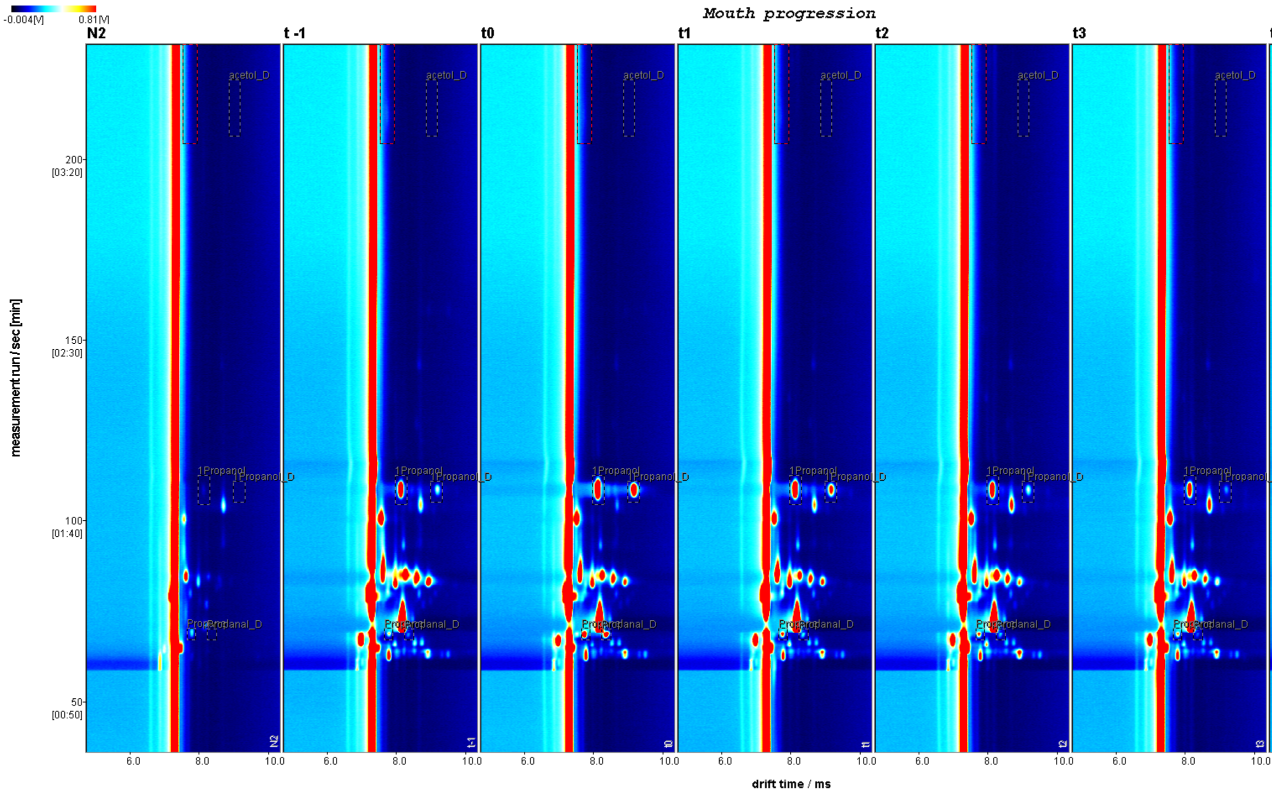


**Figure S1.** Sequence of the GC-IMS Chromatograms of the end-tidal oral exhaled air of a participant during the 1,2-PD intervention proptocol.

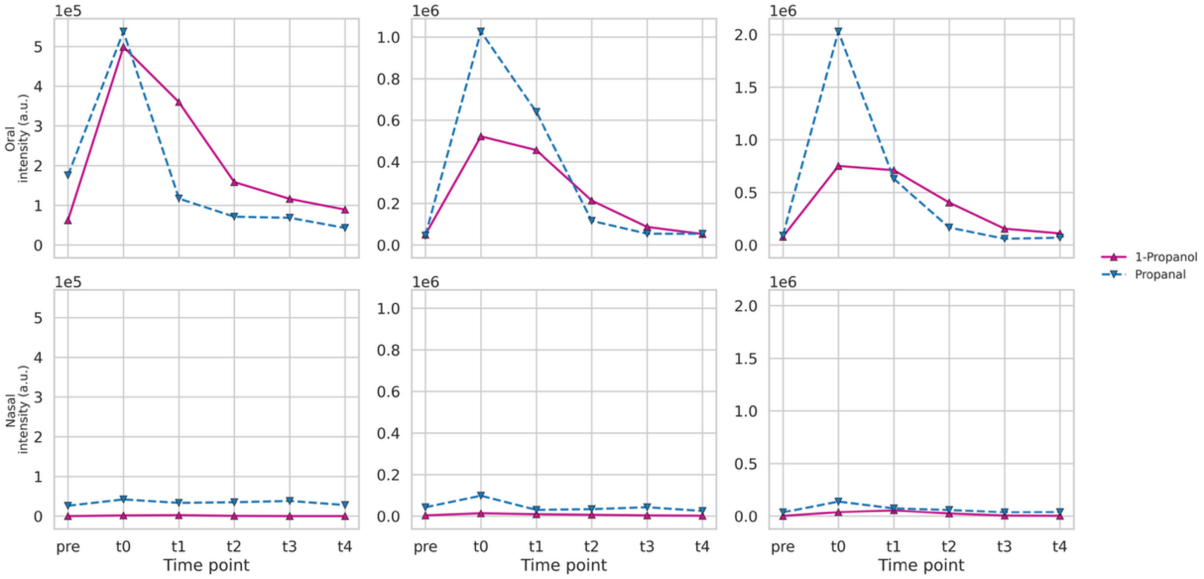


**Figure S2.** Exemplary temporal profiles showing the progression of 1-propanol and propanal through the intervention study.

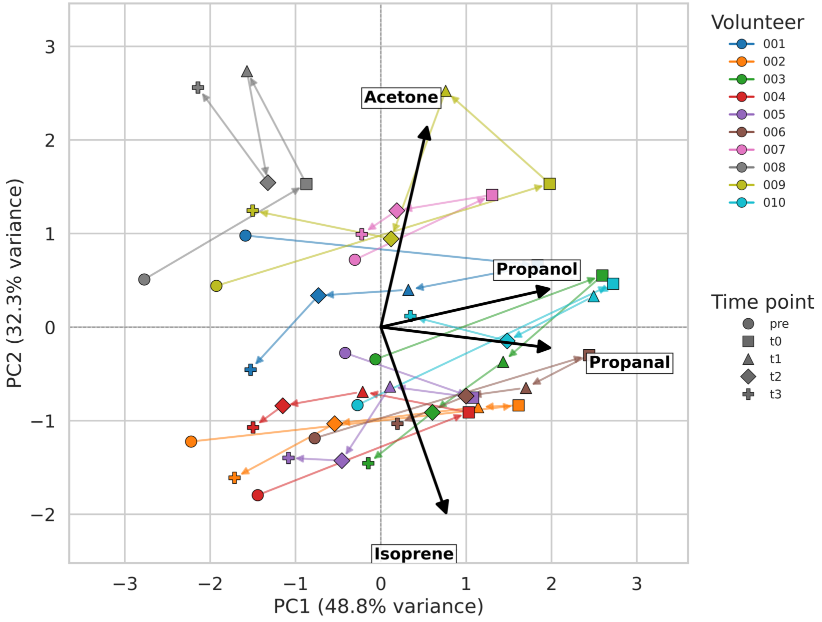


**Figure S3.** Individual trajectories through PCA space, oral samples, t_pre to t3, one path per volunteer. The marker shape encodes time point; each volunteer is represented with a different color. The 4 eigenvectors corresponding to the selected VOCs, namely propanol and 1-propanol (intervention-associated) and aceton, and isoprene (systemic compounds, unaffected by the intervention). The connected trajectory, illustrate both the shared rise-and-return pattern and the considerable inter-individual variability in its timing and magnitude.

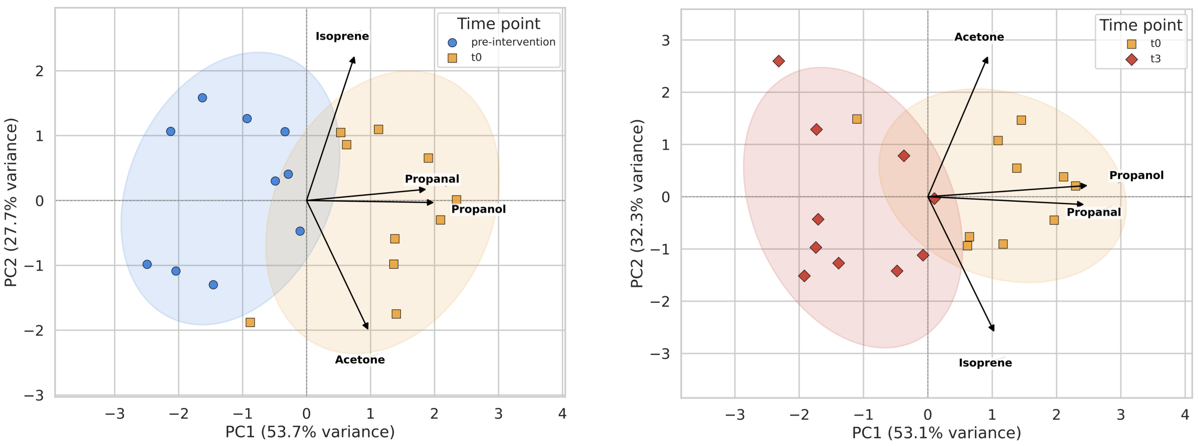


**Figure S4.** PCA biplots of oral samples, computed separately for two subsets of the sampling sequence: ($t_{pre}$ , $t_0$ ) samples subset, capturing the immediate post-intervention shift (left plot); ($t_0$ , $t_3$) samples subset, capturing its reversal during washout (right). Arrows denote variable loadings. In both panels, separation between time points occurs predominantly along the loadings for propanal and 1-propanol.
